# GABAergic circuits reflect different requirements for emergent perception in postnatal mouse neocortex

**DOI:** 10.1101/2023.11.21.568139

**Authors:** Filippo Ghezzi, Liad J. Baruchin, Ngoc T. Ha, Mark J. Shah-Ostrowski, Alessandra G. Ciancone Chama, Jacqueline A. Stacey, Simon J.B. Butt

**Affiliations:** Department of Physiology, Anatomy & Genetics, University of Oxford Oxford, OX1 3PT, UK

## Abstract

Information transfer in the mammalian cerebral cortex is dependent on locally-projecting GABAergic interneuron circuits that are widely assumed to be uniform across neocortical areas. We demonstrate that this does not hold true during the highly dynamic period of postnatal life prior to the onset of active sensory exploration. During this time, a subset of interneuron defined by expression of the neuropeptide somatostatin differentially contribute to sensory-evoked activity in primary somatosensory and visual cortices. This functional divergence between the two areas is explained by differences in the composition of somatostatin interneuron subtypes and the transient circuits formed by these cells; the somatosensory circuit representing an adaptation to control early neonatal touch information. Understanding such area-dependent differences will promote our endeavours to understand the aetiology of developmental psychiatric disorders.

**Summary Sentence:** Cortical circuits are adapted to the local information processing demands of the developing brain

---

During development, nervous systems face a significant challenge, to integrate and interpret sensory information at a time when the building blocks of the neural circuitry are being formed. During embryonic development many of the processes fundamental to nervous system development are constrained by genetic and molecular cues [1]. However, at the onset of perinatal life, physiological signals start to direct and ultimately ensure faithful representation of both external and internal states to enact appropriate behavioural responses [2, 3]. In the mammalian cerebral cortex, a number of common strategies have been identified that play out during this later period of development [4]. These include transient neuronal populations and circuits [5–7], as well as critical periods for plasticity - time points during which representation of the sensory environment is consolidated within the cytoarchitecture of cortical circuits [8]. Locally-projecting GABAergic interneurons play an important role in both and further serve to coordinate early activity, promote neuronal maturation and induce synaptogenesis [7, 9–12]. The central nature of GABAergic activity to these processes has led to several studies exploring the contribution of interneuron subtypes to early activity and revealed fundamental mechanistic insight into the interaction between GABAergic interneuron and pyramidal cells (PYRs) at early ages [9, 13, 14]. However, we still do not understand how these neurons act on the millisecond time scale to synchronize and constrain nascent activity; a key omission given that these cells likely contribute to formative events such as the dominant spontaneous activity of early neocortex – β-frequency, spindle burst oscillations [3, 15, 16]. Further, we have little understanding if any such functionality is uniform across cortical areas. This is an important consideration given differences in the timing of critical periods between primary sensory areas in neocortex [8], and the fact that early interneuron dysfunction is thought to underpin neurodevelopmental psychiatric disorders with area-specific differences having the potential to manifest as distinct pathophysiology [17].

We adopted an *in vivo* optotagging approach, previously used in adult neural circuits [18] to identify the contribution of GABAergic interneurons to both spontaneous and sensory-evoked activity in postnatal mouse neocortex on the millisecond time scale. We found that while somatostatin (SST)-expressing interneurons participate in early spontaneous activity across both primary visual (V1) and whisker somatosensory (S1BF) cortex, only those present in the latter regulate the fast sensory-response prior to the onset of active sensation. Further, a transient translaminar circuit formed by thalamo-recipient infragranular SST interneurons in somatosensory cortex [7] is absent at corresponding postnatal ages in visual cortex with corresponding differences in the composition of infragranular SST subtypes [19, 20] found between these areas. We propose that the transient SST interneuron circuit found in the somatosensory cortex represents an adaptation necessary to control early touch information in this area. Furthermore, such differences in the construction of formative circuits have significant implications for our understanding of the genetic, molecular and cellular mechanisms that instruct emergent function and ultimately, the aetiology of neurodevelopmental psychiatric disorders [17].

## SST interneurons contribute to discontinuous beta-frequency oscillations in a manner independent of sensory area

A diverse array of cortical GABAergic interneuron subtypes operate over a range of temporal and spatial scales to control regular spiking (RS), pyramidal cell function and thereby higher order cognition throughout life [21, 22]. Of these, one subtype defined by expression of SST is thought to be an important player in cortical development, instructing circuit maturation and synaptogenesis [7, 11, 12]. To understand the contribution that this interneuron subtype makes to formative neural activity we developed an optotagging approach reliant on conditional expression of ChannelRhodopsin2 (ChR2) in SST-expressing interneurons by crossing an *SST-ires-Cre* transgenic mouse with the *Ai32* reporter line (**Figure 1A**, **Figure S1A**). We further explored the wider population of GABAergic interneurons originating in the medial ganglionic eminence (MGE), that include SST interneurons as well as fast spiking (FS), parvalbumin-expressing (PV) cells, using an *Nkx2-1Cre* line [23] to drive ChR2 expression (**Figure 1A**, **Figure S1B**, **Figure S2**). Pups with these combinations of alleles allowed us to *post hoc* identify SST and Nkx2-1 single units (SU) using optotagging (**Figure S1, C-F**), and thereby determine the relative contribution of these cell types to spontaneous (**Figure 1**) and sensory-evoked *in vivo* neural activity (**Figure 2**) in both S1BF and V1 during postnatal life. Most experiments were performed under urethane (UR) anaesthesia [24, 25] to provide a uniform state across the ages tested, with a further, smaller number performed in awake (Aw), head-fixed pups at early ages only. During the initial postnatal period, prior to the onset of active sensory exploration, we observed characteristic beta (β)-frequency (10-30 Hz) activity (**Figure 1** **B** and **C**) – termed spindle bursts – in both cortical areas [15, 26]; activity that appeared more discontinuous under anaesthesia (**Figure S3, A-D**). RS cells and optotagged interneuron SUs fired almost exclusively within and were entrained by these β-frequency oscillations (**Figure 1, D, F, H**) across the depth of the cortical column at early ages. Our failure to optotag interneuron SUs at early ages in V1 precluded comparison with S1BF during the earliest time window, P5-8 (**Figure 1** **D** and **E**). At this age in S1BF, RS single units in granular L4 were more entrained than those in infragranular L5/6 (**Figure 1D**); a distinction that subsequently emerged between L2/3 and L5/6 RS SUs across both areas at P9-13 (**Figure 1, F** and **H**), consistent with the maturation of the feed-forward L4 to L2/3 circuit in advance of active sensory exploration [27]. In S1BF, Nkx2-1 and SST SUs showed similar degrees of entrainment (**Figure 1F**) but the peak phase of SST interneuron firing was significantly delayed from that of both RS and Nkx2-1 units (**Figure 1G**), pointing to discrete roles for interneuron populations in controlling pyramidal cell activity prior to the onset of active whisking. In V1, SST interneurons were highly entrained compared to Nkx2-1 SUs (**Figure 1H**) and phase shifted relative to RS but not Nkx2-1 SUs during β-frequency oscillations (**Figure 1I**). Thus, it appears that SST interneurons contribute to spontaneous activity in V1 prior to eye opening, more so than the broader class of Nkx2-1 interneurons that includes presumptive FS, PV-expressing cells.

**Figure 1.**
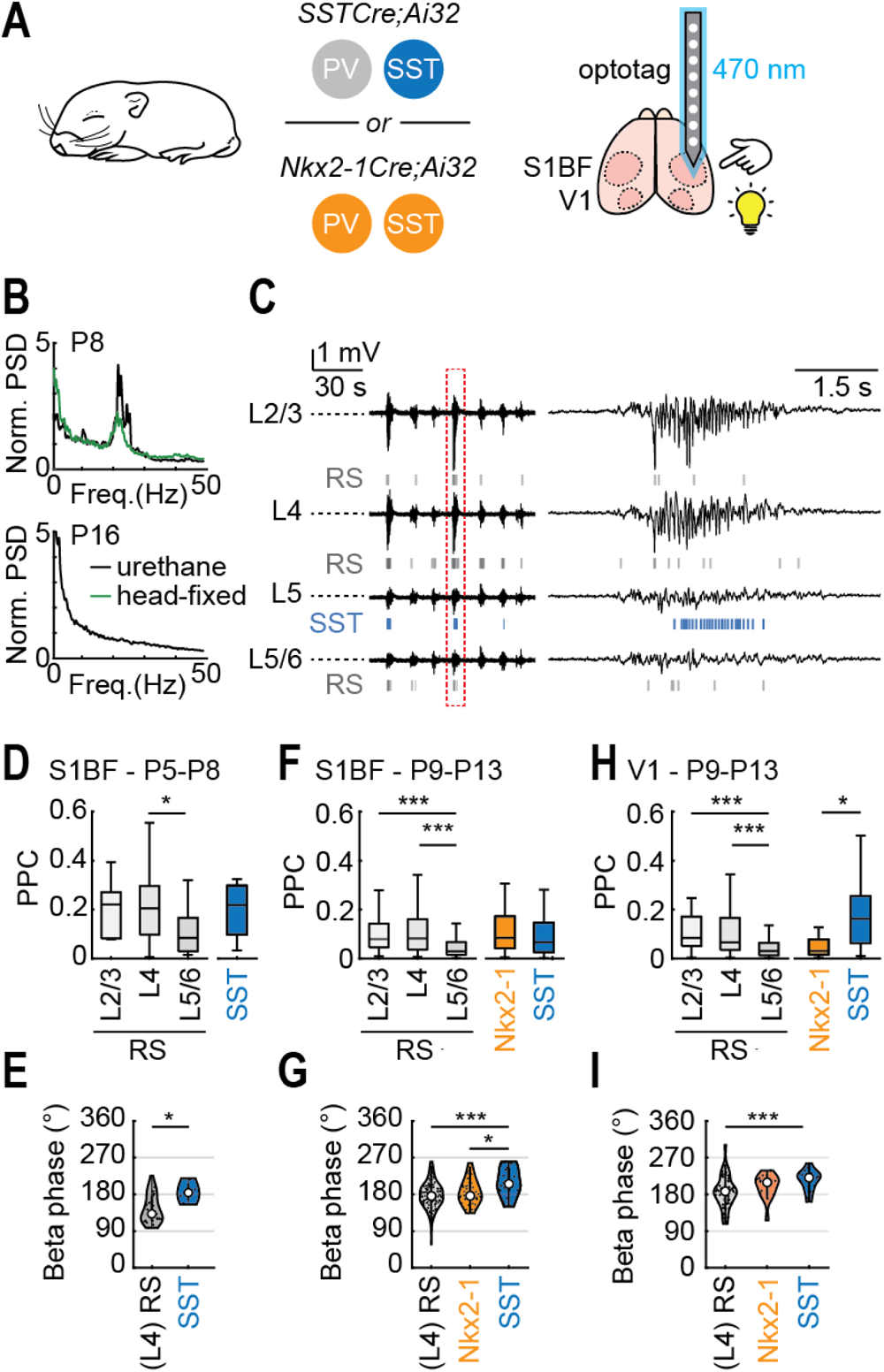
SST interneurons are recruited during early postnatal spontaneous activity in both S1BF and V1. (A) Genetic strategies used to optotag and identify SST (blue) or SST and PV (orange) interneurons in both somatosensory (S1BF) and visual (V1) areas of mouse neocortex (right) *in vivo*. (B) Power spectra for spontaneous activity recording from V1 at P8 (top) and P16 (bottom). (C) Spontaneous *in vivo* activity with the β-frequency spindle burst in the red dashed box shown on an expanded time scale to the right. RS, the firing patterns of regular spiking single units; the discharge of an SST single unit is shown in blue. (D) Pairwise phase-consistency (PPC) values between spike times and β-frequency oscillations and (E) average firing phase for RS and SST single units during β oscillations in S1BF between P5-8. (F,G) Corresponding data for S1BF between P9-13, just prior to active whisking, and (H,I) in V1 for P9-13, prior to eye opening; SST data shown in blue; Nkx2-1 in orange. * P < 0.05; *** P < 0.001.

**Figure 2.**
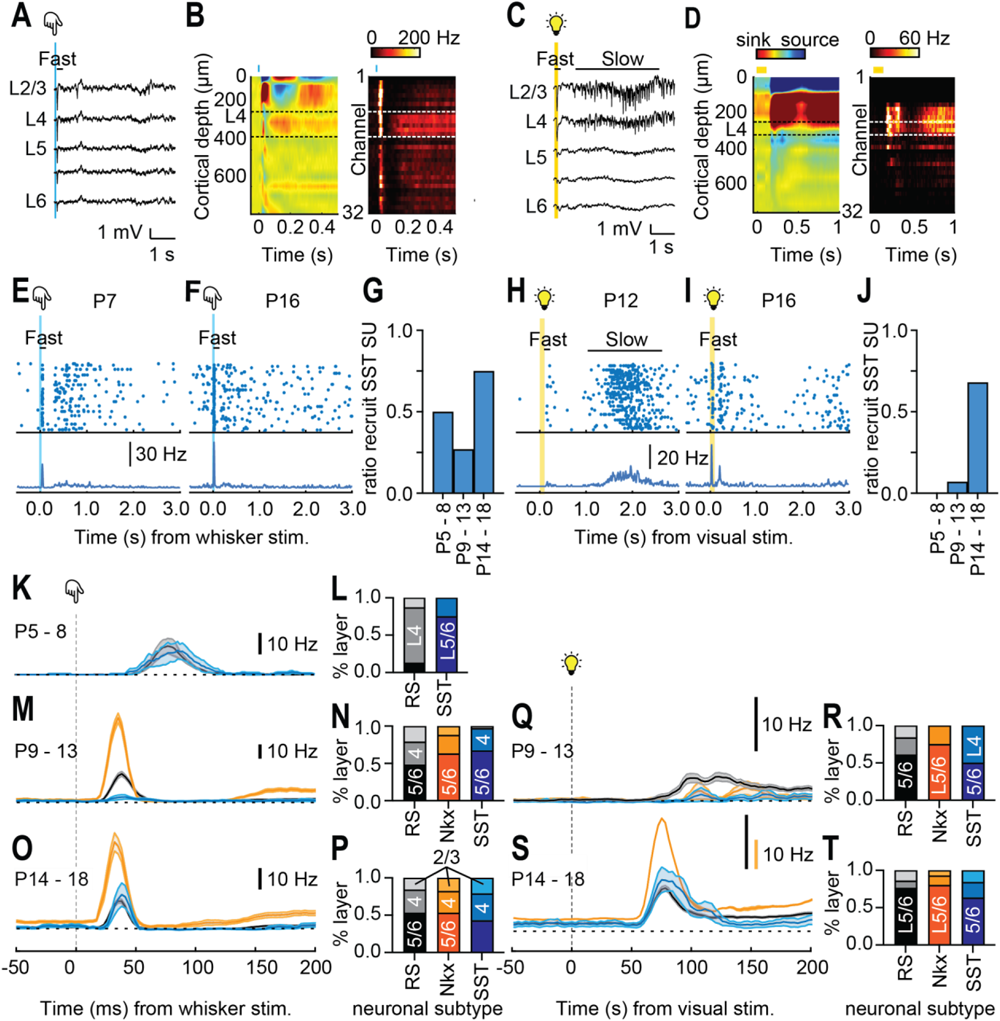
SST interneurons are recruited during the short latency sensory-evoked response in early postnatal S1BF but not V1. (**A**) Multi-whisker stimulation (onset indicated by the blue line) triggers a fast response across the depth of cortex in S1BF with corresponding (**B**)(left) current source density (CSD) plot and (right) multiunit activity (MUA). (**C**) From P8 onward, a100 ms flash of white light (yellow bar) can trigger a sensory response in V1. Prior to eye opening this consists of both fast (<200 ms from stimulus onset) and slow components as seen in (**D**) the CSD and MUA traces. Raster plot (top) and peri-stimulus time histogram (PSTH) for optotagged (**E**) P7 and (**F**) P16 SST interneurons in S1BF. (**G**) Ratio of recorded SST SUs recruited in S1BF over the period of development studied. Corresponding data for (**H-J**) SST interneurons recorded in V1. (**K**) Grand-averaged PSTHs for RS (grey) and SST (blue) neurons in P5-8 S1BF. (**L**) layer location of whisker stimulus-responsive units. Corresponding data for (**M,N**) P9-13 (**O,P**) P14-18 S1BF including Nkx2-1 SUs (orange). (**Q,R**) Same as for (**M,N**) and (**S,T**) same as for (**O,P**) but for V1 SUs.

## Sensory recruitment of SST interneurons differs between postnatal S1BF and V1

Short latency, sensory-evoked responses can be elicited in mouse S1BF during the first postnatal week (Figure 2**, A** and **B**) [25] but are delayed in V1 until ∼P8, at which time point responses can be recorded in response to a flash of light through the closed eyelids [26, 28]. Visual responses evoked prior to eye opening (defined here as P14) consist of a fast response (<200 ms from stimulus onset) (Figure 2C) that spans the depth of the cortical column followed by a prolonged, slow response with associated multi-unit activity (MUA) reverberating in L2-4 for over 1 s (Figure 2D). As previously reported [24, 25], anaesthesia had no impact on sensory-evoked except for increased numbers of SUs recruited during the slow response in V1 (**Figure S3 E-G**). In both areas, fast-sensory responses were underpinned by activity in RS SUs which were recruited at short latency to the stimulus in increasing number as development progressed (**Figure S4, A-D**). The recruitment of SST SUs however varied over development and between areas (Figure 2**, E-J**). In S1BF, we observed sensory-driven SST units across all the time windows examined (Figure 2**, E** and **F**) albeit that there was a dip in the proportion (Figure 2G) and strength (Figure 2M) of SST interneuron recruitment just prior to the onset of active whisking (P9-13). In V1, most SST units only contributed to the slow response prior to eye-opening (Figure 2H) but were recruited abruptly during the fast response post-eye opening (Figure 2**, I** and **J**). The same pattern was observed for Nkx2-1 units in V1 (**Figure S4, F-H**), whereas in S1BF Nkx2-1 units were recruited both before and after the onset of active whisking (**Figure S4E**). Examination of the short latency sensory responses across early development in S1BF (**Figure 2, K-P**; **Figure S4, I-K**) and V1 (Figure 2**, Q-T**; **Figure S4, L** and **M**) revealed that in the former, SST neuron – predominantly located in L5/6 (Figure 2L) - fire coincident with RS units in L4 (**Figure2 K** and **L**; **Figure S4I**) during the earliest time window recorded. Subsequently, feed-forward, short-latency recruitment of Nkx2-1 neurons becomes the dominant feature with SST SUs activated at a delay. The switch in the latter is consistent with reported the timing of the transient thalamic input onto S1BF SST neurons [12] with the overall pattern of neuronal recruitment matching that reported in adults [29] post-P9. In V1, the few fast-responsive interneurons observed were recruited at a delay to RS units prior to P14 (**Figure 2** **Q and R;** **Figure S4L**), but rapidly switched to an adult-like configuration immediately after eye opening (Figure 2S and **Figure S4M**). These data demonstrate that in S1BF, SST and Nkx2-1 interneurons contribute to early, fast sensory-driven information transfer prior to active sensory exploration, which is not the case in V1. However, both sensory areas exhibit the same adult-like pattern of interneuron recruitment following the onset of active sensation (Figure 2 **O and S**).

## Difference in SST recruitment are underpinned by circuit and subtype differences between S1BF and V1

Our *in vivo* optotagging data point to different developmental trajectories in postnatal GABAergic circuitry between the two primary sensory areas studied. In S1BF, thalamo-recipient L5/6 SST interneurons sequentially innervate cortical layers in an inside-out manner during the first two weeks of life [30, 31] and include – during the critical period for plasticity for L4 – a reciprocal synaptic loop with spiny stellate neurons (SSNs) (Figure 3A)[7]; a transient connection that predates the onset of fast feed-forward inhibition mediated by local parvalbumin (PV)-expressing interneurons[32, 33]. To explore the corresponding circuitry in postnatal V1, we mapped total columnar GABAergic synaptic input onto pyramidal cells (PYRs) *ex vivo* using laser scanning photostimulation (LSPS) of caged glutamate. Recorded L4 neurons were voltage-clamped at the reversal potential for glutamate (E_Glut_) to observe LSPS-evoked IPSCs. Across all time points tested, both pre-and post-eye opening, we found no evidence of prominent translaminar GABAergic connections onto L4 cells in this area, instead input was largely confined to the immediate layer (Figure 3B). We further confirmed the absence of the SST-mediated translaminar connections in V1 using a Cre-dependent LSPS approach to map the presynaptic location of afferent SST interneurons [31]. The resultant SST-specific input profiles matched those obtained for total GABAergic input albeit shifted towards the L4/L5 boundary at P9-13 (Figure 3C).

**Figure 3.**
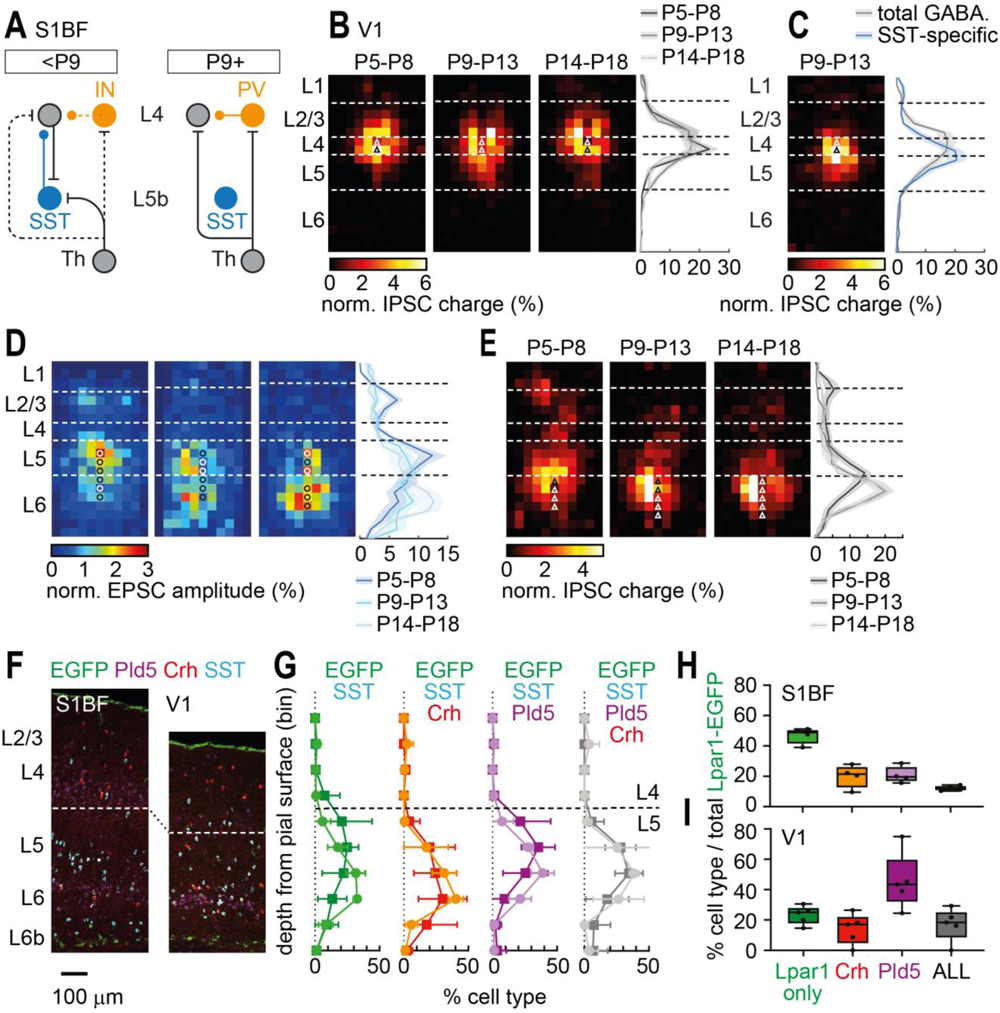
Postnatal GABAergic circuits in V1 are distinct from those observed in S1BF. (**A**) Schematic of the transient L5b-L4 synaptic circuit present in S1BF before the onset of feed-forward inhibition at P9 (*7*). (**B**) LSPS-derived maps of GABAergic input onto L4 PYRs in V1 across 3 postnatal time windows; (right) plot showing normalised input profiles across development. (**C**) SST interneuron specific input map at P9-13; (right) SST-specific profile (blue line) plotted against total GABAergic input (grey). Maps of (**D**) glutamatergic input onto SST interneurons and (**E**) GABAergic input onto PYRs, both in L5/6 of V1. (**F**) RNAscope *in situ* hybridisation for SST interneuron markers. Dashed line, L4/L5 boundary. (**G**) Average distribution of Lpar1-EGFP interneurons subtypes across depth of S1BF (light colours) and V1 (dark colours). Proportion of Lpar1-EGFP subtypes in (H) S1BF and (I) V1. 2-way Anova for difference in cell types according to cortical area (S1BF *vs.* V1): F(3, 28) = 10.20, P<0.001.

The feed-forward input from L5b SST interneurons onto L4 SSNs observed in S1BF precedes the onset of local, feed-forward inhibition mediated by PV+ basket cells [32, 33](Figure 3A). This led us to question the role of V1 SST cells located at the L4/5 border at early ages, given that they do not participate in the fast, sensory response (Figure 2H and J). We used an *ex vivo* optogenetic approach, first examining thalamic innervation and the onset of fast feed-forward inhibition in V1 (**Figure S5A**). This revealed that these SST interneurons innervating L4 PYRs – in contrast to those in L5b S1BF [7, 12], do not receive direct thalamic input (**Figure S5, B-E**) with the onset of fast, feed-forward inhibition delayed in V1, coincident with eye opening (**Figure S5, G-I**). We then generated *Lhx6-EGFP;SST-Cre;Ai32* pups to examine interactions between SST, putative-PV (Lhx6+ but not ChR2-responsive) interneurons and PYRs; *Lhx6* being a transcription factor downstream of *Nkx2-1* [34]. PV cells received increased innervation from SST interneurons independent of cortical area in the immediate time window prior to the onset of active sensation (**Figure S6**), however this originated from L5 in S1BF (**Figure S6, A**) but was local to L4 in V1 (**Figure S6, C**). Further, the amplitude of IPSCs elicited from SST interneurons onto L4 SSNs and PV cells was similar in S1BF (**Figure S6, B)**, whereas in visual cortex, SST synaptic inputs onto PV interneurons were significantly larger than those observed in PYRs (**Figure S6, D**). Finally, to unequivocally establish the presence of distinct early SST interneuron circuits in postnatal V1, we returned to LSPS of caged glutamate in *SSTCre;Ai9* pups and tested for translaminar excitatory connections onto infragranular SST interneurons akin to that observed in postnatal S1BF (Figure 3A)[7]. While the L4 excitatory input was absent – identifying a further difference to S1BF, our LSPS maps revealed that V1 SST cells received translaminar input from supragranular L2/3 PYRs during the earliest time window (P5-8) (Figure 3D). This observation caused us to revisit and extend our mapping of total GABAergic input onto V1 PYRs beyond L4 through postnatal development. Our survey revealed no reciprocal GABAergic connection from infragranular layers onto L2/3 PYRs (**Figure S7, A** **and B**) but did identify a subset of infragranular PYRs that receive translaminar GABAergic input from L2/3 during the earliest time window recorded (Figure 3E; **Figure S7, C** **and D**). The presence of these parallel glutamatergic and GABAergic translaminar connections (Figure 3**, D and E**) suggests an instructive role for L2/3 in the co-ordination of early infragranular activity in V1 that is quite distinct from the model observed in postnatal S1BF (Figure 3A). Further, infragranular SST interneurons in V1 do not sequentially innervate cortical layers in an inside-out manner as found in S1BF (Figure 3C; **Figure S7**) [7]. This suggests that either infragranular V1 is populated with SST interneuron subtypes distinct from those found in S1BF or that the same subtype fulfils different roles across sensory areas during early postnatal development. *Lpar1-EGFP* (EGFP) SST interneurons – the subtype responsible for the transient translaminar connection in S1BF [7], were found in similar numbers in infragranular layers of both primary sensory areas during the early postnatal period (**Figure S8**). Fluorescent *in situ* hybridisation for transcriptomic markers of infragranular SST interneurons revealed that two markers, *Pld5* and *Crh* [19], delineated *Lpar1-EGFP* subtypes with similar distributions (Figure 3**, F** and **G**) but differing proportions across both primary sensory areas. The *Lpar1-EGFP* only subtype was more prevalent in S1BF (Figure 3H) whereas EGFP cells that co-expressed *Pld5* were more evident in V1 (Figure 3I). These data show that relatively small differences in the composition of interneuron subtypes between these two primary sensory areas – akin to that reported in adult mice [35–37] – can result in significant differences in early GABAergic circuitry and recruitment during formative sensory activity.

## SST interneurons provide feed-back control in V1 but transiently regulate sensory input in a feed-forward manner in postnatal S1BF

Differences in both the recruitment and identity of SST interneurons point to a different role for this interneuron class during early postnatal life across the two sensory areas studied. To establish if this is the case, we pursued both opto-and chemogenetic approaches to understand how these cells regulate early sensory activity. Optogenetic activation of SST interneurons (via conditional expression of ChR2 previously used for optotagging) coincident with the sensory stimulus typically attenuated RS single unit activity both pre-and post-active sensory exploration across both V1 (**Figure4**, **A and C**; Figure S9, **D and E**) and S1BF (Figure 4**, B;** **Figure S9**, **B and C**) – consistent with a net inhibitory effect – except for the P9-13 window in S1BF when we observed an increase in disinhibited RS units (**Figure S9, A-C**). This likely reflects the non-physiological nature of our opsin-mediated stimulation, and we believe, is unlikely to be a prominent signalling motif due to the limited recruitment of SST interneurons during this time window (**Figure2, G** and **M**) under the conditions examined here. We then used a chemogenetic approach [38] to examine the impact of a reduced SST interneuron activity on sensory-evoked responses in both areas. Injection of the DREADD viral vector *AAV8-KORD-IRES-mCitrine* into newborn *SSTCre;Ai9(tdTomato)* animals resulted in expression of KORD in SST interneurons (**Figure S10, A-C**) and a reversible, drug-dependent (Salvinorin B; SalB) hyperpolarisation of transfected SST interneurons recorded *ex vivo* P7-9 (**Figure S10, D-G**). For *in vivo* experiments, baseline spontaneous and sensory-evoked activity was recorded prior to injection of SalB in pups with control SST^GFP^ (transduced with an AAV-GFP virus) or SST^KORD^ interneurons (Figure 4**, D-J**) (**Figure S11**). SalB injection in control GFP animals (**Figure S11, A** and **B**) or injection of vehicle in KORD-injected animals had no effect on activity but in SST^KORD^ pups we observed different effects depending on the primary sensory area. In S1BF, acute exposure to the drug SalB resulted in an increase in MUA – in contrast to prior observations that used chronic silencing of SST cells [39] – but only during the P5-8 time window (Figure 4**, D** and **E**) (**Figure S11, C-E**). In V1, injection of SalB had no effect on the sensory evoked MUA at either of the ages tested (Figure 4**, E** and **F**) (**Figure S11, F-H**), but led to an increase in spontaneous, background SU activity (Figure 4**, H** and **I**) accompanied by a corresponding decrease in the percentage of spikes occurring within β-frequency, spindle burst activity prior to eye-opening (Figure 4J). These data confirm that SST interneurons directly regulate early sensory information transfer in S1BF in a feed-forward manner [7]. This role is distinct from that seen in V1, where SST interneurons are confined to a feed-back role controlling baseline activity and spread of sensory activity through early life [40].

**Figure 4.**
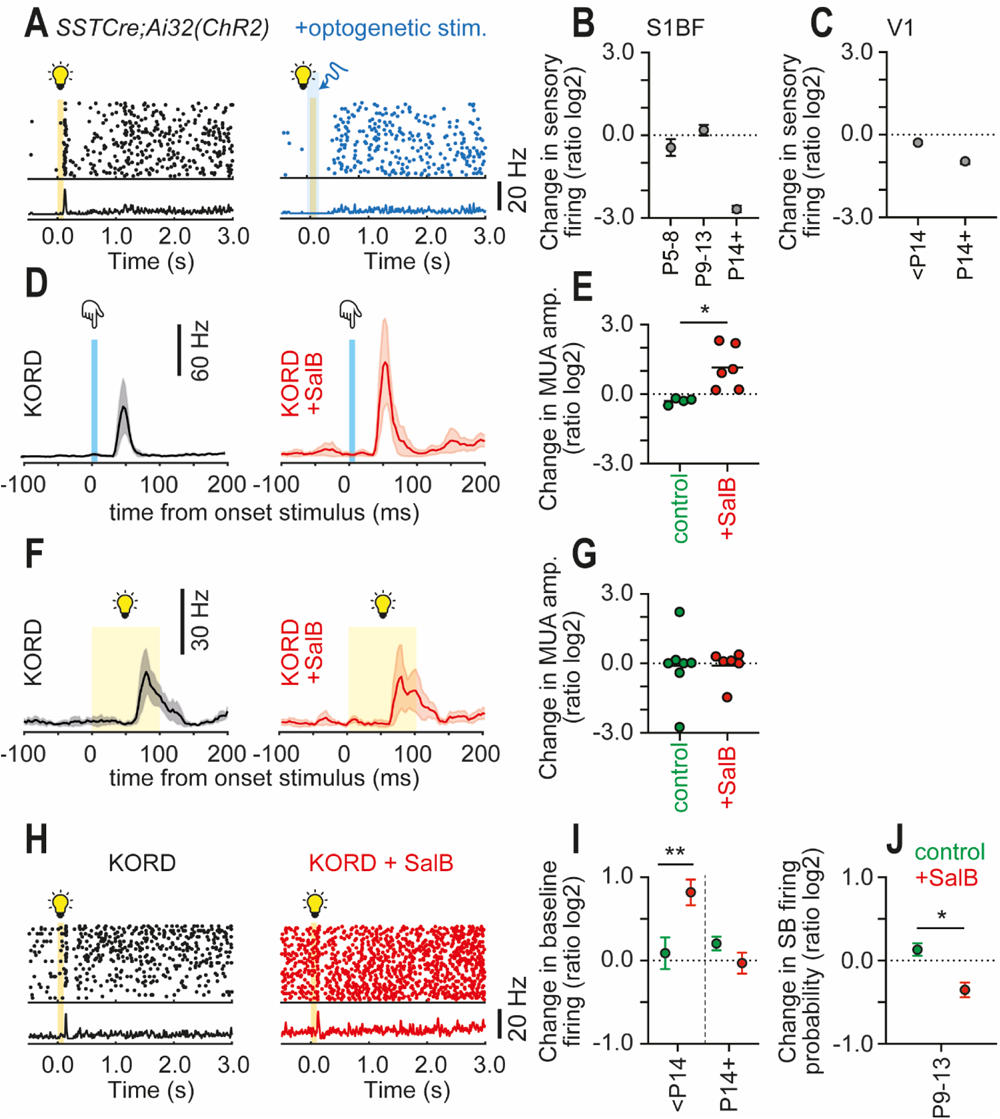
SST interneurons regulate the short latency sensory response in postnatal S1BF but not in V1. (**A**) Impact of simultaneous ChR2 optogenetic stimulation of SST interneurons on the sensory response of a representative RS single unit in V1. Average effect of ChR2-mediated SST interneurons activation on short latency sensory responses recorded in (**B**) S1BF and (**C**) V1 over postnatal development. (**D, E**) Effect of bath application of Salvinorin B (SalB) on the sensory evoked response in S1BF in P5-8 animals in which the K-opioid receptor (KORD) was conditionally-expressed in SST interneurons. Welch’s t-test control *vs.* +SalB, *p = 0.012 (**F, G**) Corresponding data for V1. 2-tailed t-test control *vs.* SalB, p = 0.987. (**H**) Raster plot showing altered RS SU activity in V1 following SalB perfusion, with (**I**) an increase in baseline firing observed prior to eye opening (control vs. +SalB at <P14, Wilcoxon rank sum test: Z = 2.6, **p = 0.01; control vs. +SalB at P14-P18, Wilcoxon rank sum test: Z = −0.4, p = 0.69.) with (**J**) a parallel decrease in probability of firing within beta-frequency, spindle bursts (two sample t-test: t(79) = −2.5, *p= 0.02).

## Discussion

The sum of these data suggest that the early GABAergic circuits are adapted to the information processing demands of a given sensory area during early postnatal life. More specifically, SST interneuron subtypes in S1BF receive direct thalamic input to regulate sensory responses within the first postnatal week and drive circuit maturation [7, 11, 12]. For V1, there are no such concerns as light-evoked responses are not present until early in the second postnatal week (P8)[26, 28] and therefore no need for SST interneurons to engage in resource-expensive transient networks, but rather they can simply adopt a feed-back role, regulating synchronised spontaneous activity in the cortex [40], in line with their eventual adult function. We speculate that feed-forward engagement of SST interneurons evolved in early postnatal S1BF because of the need to control touch sensory responses during the perinatal period [2]. SST interneurons are ideally placed to perform such a function given their early birthdate [41], preponderance in deep layers, and morphological attributes including ascending axons that ramify through the more superficial, later-born cortical layers [7]. Moreover, their role in development is multifaceted, extending beyond immediate control of information transfer as shown here, to include synaptogenesis [11] and neuronal maturation [12]. It remains to be seen if this specialisation extends to other sensorimotor areas but we suggest that it is unlikely to be found in auditory and association areas whose later developmental trajectory is similar to that observed in V1; a finding that would be consistent with the reported similarity in interneuron subtypes previously reported across anterior-motor areas and V1 [20]. Our results parallel other recent reports of differences in adult interneuron circuits across cortical areas [35–37], extend this to *in vivo* function, and further demonstrate that a small shift in the make-up of GABAergic interneuron subtypes can have profound consequences for the regulation of neural activity and resultant information transfer from early on in life; an observation that raises fundamental questions about our pursuit of GABAergic interneuron-related genetic and molecular targets in our understanding of neurodevelopmental psychiatric disorders [17].

## Acknowledgments

We would like to thank our funders as well as current and former members of the Butt lab who have contributed to discussions.

## Author contributions

Conceptualization: SJBB; Methodology: FG, LJB, JAS and SJBB; Investigation: FG, LJB, NTH, MS-O, AGCC, JAS, SJBB; Visualization: FG, LJB, SJBB; Funding acquisition: FG, LJB, SJBB; Project administration: SJBB; Supervision: FG, LJB, SJBB; Writing – original draft: FG, LJB, SJBB; Writing – review & editing: FG, LJB, NTH, MS-O, AGCC, JAS, SJBB

## Data and materials availability

All data, code, and materials used in the analysis are available by contacting the corresponding author. Datasets will be released via ora.ox.ac.uk on publication.

## SUPPLEMENTARY MATERIALS

### Materials and Methods

#### Animal models

All experiments were approved by the University of Oxford ethical review committee and performed in accordance with Home Office project license P861F9BB7 under the UK Animals (Scientific Procedures) 1986 Act. The following mouse lines were used *SST-ires-Cre* (Ssttm2.1(cre)Zjh/J), *Lhx6-eGFP* (Tg(Lhx6-EGFP)BP221Gsat), *Ai32* (Gt(ROSA)26Sortm32 (CAG-COP4*H134R/EYFP)Hze/J), *Ai9* (Cg-Gt(ROSA)26Sortm9(CAG-tdTomato)Hze/J), *Lpar1-EGFP* (Tg(Lpar1-EGFP)GX193Gsat), *Nkx2-1-Cre* (Tg(Nkx2-1-cre)2Sand/J), *floxed-stop-P2X2R* (R26::P2x2r-EGFP), and *Olig3-Cre* (Olig3tm1(cre)Ynka). Neonatal mice of either sex – with the exception of the *Lpar1-EGFP* transgenic which is Y chromosome linked - were generated via targeted breeding and used for experiments between postnatal day (P)5 and P18.

#### *In vivo* electrophysiology

*In vivo* electrophysiology experiments were performed on neonatal mice either under urethane anaesthesia (0.5–1.0 g/kg) or awake head-fixed. For the former, as soon as animals lost their tail and pedal reflexes, they were head-fixed in a stereotaxic apparatus and a small craniotomy was opened over S1BF (2.7–3.2 mm lateral and 2.4–2.8 mm rostral to lambda) or V1 (2.2–2.5 mm lateral and 0.5–0.7 mm rostral to lambda). For awake head-fixed experiments, mice were anaesthetised with isoflurane (in O_2_; 5% for induction, 2-3% for maintenance) and placed on a heating pad. The skin over the skull was then removed, the skull cleaned with 3% H_2_0_2_ and a head-fixation apparatus was made, using dental cement and two light plastic tubes. Finally, a craniotomy was opened over S1/V1, and the animal was head-fixed in the same stereotaxic apparatus and allowed to recover from isoflurane anaesthesia. Within the stereotaxic apparatus, mice were placed on a heating pad to provide thermoregulation. Recordings were performed with a single shank, 32-channel multi-electrode array (A1x32-Poly2–5mm-50s-177 or A1x32-Poly3-10mm-50-177; NeuroNexus, USA) or corresponding optoelectrodes, slowly inserted (∼1 µm/s) perpendicular to the brain surface at a depth of ∼800 µm using a micromanipulator (SOLO-50, Sutter Instruments, US). After 30 mins. interval to allow for tissue recovery, data were acquired at 20 or 30 kHz with an OpenEphys acquisition board. Recordings lasted for ∼ 60 mins and consisted of a 20-min baseline recording followed by optogenetic tagging and sensory stimulation sessions.

Visual stimuli (100 ms duration) were administered to the contralateral eye using a white LED light stimulus (C503C-WAN, Cree, UK) every 15-30 seconds controlled via an input-output board (National Instruments, UK). Whisker stimulation was obtained by inserting the contralateral whiskers into a glass cannula connected to a piezoelectric unit (Thorlabs, US) controlled by piezoelectric amplifier (E-650 Piezo Amplifier; PI; Germany). Either a single stimulus or pair stimuli (500 ms inter-stimulus interval) were applied every 15-30 s.

Optogenetic spike-tagging was performed through a fibre-coupled blue laser (Stradus 473/70mW; Laser 2000, UK), positioned close to the craniotomy or connected to an optoelectrode. Long duration (50 ms) laser pulses were employed to account for reduced excitability of ChR2 during early postnatal development. One optogenetic session, characterized by 100 laser pulses delivered at 0.1 Hz frequency, was conducted immediately before the sensory stimulation session. Isolated single units (SU) were considered optotagged interneurons if they significantly increased firing rate during a 470 nm LED light pulse (**Figure S1, C** and **D**) compared to background activity. Out of a total of 2113 SU obtained from *SSTCre;Ai32* animals, 109 (5%) were tagged as SST interneurons while 110 out of 875 SU (13%) recorded from *Nkx2-1Cre;Ai32* animals were tagged as Nkx2-1 interneurons. We were unable to optotag interneuron units prior to P7 in V1, likely due to the immature electrophysiological properties of both SST and Nkx2-1 interneurons at this time. Prior to P14, SU invariably exhibited slow spike waveforms (**Figure S1, C**) with interneurons only isolated from other SU and multiunit activity (MUA) based on optotagging (**Figure S1, E**). Only after the onset of active sensory exploration (active whisking/eye opening; P14 onward) was it possible to separate putative regular spiking (RS) and fast spiking (FS) single units based on the spike trough to peak latency histogram alone (**Figure S1, F**) independent of our optotagging approach. During this later time window, FS units were captured using our optotagging approach in *Nkx2-1Cre;Ai32* animals however the percentage FS optotagged SU differed across the two cortical areas with 15 of 29 FS (52%) SU optotagged in S1BF but only 9 of 55 (16%) in V1. No FS units were optotagged in *SST-ires-Cre;Ai32* animals. The distribution of optotagged SST and Nkx2-1 SU was biased toward infragranular (L5/6) layers throughout early postnatal development (**Figure S1, G-J**) - irrespective of cortical area - in line with their reported distribution.

Automatic spike sorting was performed with Kilosort2 [1], and manually refined in Phy2 [2] to classify single unit (SU) activity from multi-unit activity (MUA) and noise clusters. Generally, the minimum criteria for a cluster to be considered a SU was to be characterized by less than 0.2% of refractory period violations (±1 ms). Spike waveforms were extracted from the raw data through a customized Phy plugin and waveform half-width and trough to peak latency were measured. Sus – not distinguished based on optotagging – were further classified into regular-spiking (RS) or fast-spiking (FS) between P14-18 based on an arbitrary spike trough to peak latency cut-off value of 0.75 ms, with units falling below this number classified as FS units.

Analysis of *in vivo* electrophysiology data was performed using customized MATLAB scripts. Local field potential (LFP) was obtained by low-pass filtering (150 Hz) the raw signal. The initial 20-min baseline recording was used to extract features of spontaneous activity include β oscillation spindle burst (SB) activity at early ages (<P14). The power spectral density (PSD) of the data (between 1–50 Hz) was obtained from the entirety of the 20-min recording session using the Welch’s method. Raw PSD was normalized by multiplying each value for the corresponding frequency and dividing by the mean PSD power. Spontaneous MUA or SU activity was obtained as the average number of spikes fired over the course of the 20-min recording session.

For spike-LFP coupling, LFP signal was bandpass filtered (10-30 Hz), and the phase of the β oscillation estimated using the Hilbert transform. Phase values at the time of each SU spike were collected and a statistical test was used to evaluate whether they occurred in a specific phase of the oscillation (Rayleigh’s test for nonuniformity; [3]). Strength of the spike-LFP coupling for SU was measured by calculating a pairwise phase-consistency (PPC) value as previously reported [4] which is more robust that classical vector length method when dealing with small number of spikes[5, 6].

SBs were detected during the 20-min spontaneous activity recording in the best L4 channel (see below) as previously described [7]. In brief, raw LFP signal was bandpass filtered in the β frequency (10-30 Hz), and the envelope of the signal was estimated as the real part of its Hilbert transform. SBs events were defined as portions of the envelope signal larger than 2 times its SD, characterized by at least 3 cycles in the filtered LFP signal, and with a duration longer than 100 ms. SB event onset, offset and duration were obtained and used to evaluate entrainment of SU activity with SB events. SU spike probability within SB events was obtained as the ratio between the number of spikes occurring within predefined events over the total number of spikes fired over the 20-min baseline recording. To evaluate the statistical significance of SU spike firing within SB events, inter-spike intervals (ISIs) for each SU were shuffled 1.000 times and the probability of spike to occur within SB events calculated as before. A SU was deemed entrained by SBs if the real spike probability was larger than the shuffled probability in at least 99% of cases.

For the analysis of responses to sensory stimulation, LFP signals were first used to build a current-source density (CSD) plot. L4 was identified as the earliest sink following the sensory stimulus (see Baruchin et al., 2021 for details). Amplitude and latency of the LFP deflection in every channel were measured and the best L4 channel was selected as the channel within L4 with the largest negative LFP deflection. Each SU was then assigned to a cortical layer (L2/3, L4 or L5/6) and their peri-stimulus time histogram (PSTH) following visual stimulation obtained (bin size of 3 ms, smoothing of 15 ms) from the trial-wise spike times (raster plots). Spike times in the raster plots were also used to evaluate whether a SU was responsive to sensory stimulation, by comparing total number of spikes occurring within response windows to number of spikes fired during baseline. For the fast phase of the visual stimulation (0-200 ms after stimulus onset), two response windows were defined (0-100 ms and 100-200 ms) and compared with a window of equal length immediately before the stimulus (−100 ms-0). An analysis of variance (ANOVA) or Kruskal-Wallis test (if the spike number data were not normally distributed) on the trial-by-trial number of spikes within these windows was used to determine whether a SU was responsive during the fast phase of the visual stimulation (with p-value, p ≤ 0.05). SU spiking reliability during this phase was estimated with the Fano factor, defined as the ratio between the variance and the mean number of spikes in all trials. For the slow phase of the visual stimulation (400-4000 ms from stimulus onset), the total number of spikes occurring within this window was compared with a baseline window of equal length immediately preceding the stimulus onset; a Wilcoxon signed rank test was used to evaluate whether each SU was responsive during the slow phase of the visual stimulation. Peak SU activity (for every SUs) and MUA (for every channel) during the fast phase of the visual stimulation were stored and used for further comparison; similarly, peak MUA or average SU activity during the slow phase of the visual stimulation were measured. Responsive units to whisker stimulation were evaluated in a 100 ms window following each whisker stimulus and compared with an equal length baseline window immediately preceding the stimulus. Similar statistical methods were employed to evaluate optogenetically-tagged SU, by comparing the trial-by-trial total number of spikes during the optogenetic stimulus (50 ms) and spike number in a baseline window before the stimulus. Additionally, to be classified as optically tagged SU, they had to fire at least one spike during the optogenetic stimulus in at least 20% of the trials.

For optogenetic perturbation experiments, SU firing change during the first 100 ms of the stimulus was measured as the log2 of the ratio between firing rate during this window and a baseline window of 100 ms before the stimulus. Similarly, SU firing change during the rebound from optogenetic stimulation was obtained as the log2 of the ratio between the firing in the 100 ms following stimulus offset and firing rate during a baseline window.

#### Chemogenetic perturbation of SST interneurons in neonates

For chemogenetic experiments, P0 heterozygous *SST-ires-Cre;Ai9* pups were briefly separated from the dam and anesthetized with isoflurane. A small volume of virus (50 nL x 3 cycles; rate: 0.23 nL/s; delay: 3 s) was injected (Nanoject III; Drummond, UK) in the right S1BF (1.4 mm anterior and 1.4 mm lateral to lambda, depth of 0.4-0.7 mm) or V1 (0.1 mm posterior and 1.5 mm lateral to lambda, depth of 0.4-0.7 mm). The following viruses were used: AAV8-KORD (pAAV-hSyn-dF-HA-KORD-IRES-mCitrine; Addgene, US: 65417-AAV8) and AAV1-GFP (pAAV-CAG-GFP; Addgene: 37825-AAV1.T).

Injected mice were used for *in vivo* electrophysiological recordings as described above. An initial 20 min. control baseline was followed by sensory stimulation (50 trials). After that, Salvinorin B (SalB; Tocris, UK. 1 mg/kg, 0.1 mg/ml in saline with 1% DMSO) was injected intraperitoneally (i.p.) to activate KORD. After 20 min, a test baseline recording was obtained followed by sensory stimulation. For control experiments, either SalB solution was i.p. injected in animals previously infected with a GFP virus or saline (with 1% DMSO) was injected in animals previously infected with the KORD virus. At the end of the recording, mice were anesthetized with isoflurane (4% in O_2_) and their brain rapidly dissected and fixed using 4% PFA in PBS for 24-72h at 4°C. All brains were sliced into 50 µm-thick slices, mounted on histology slides, and imaged at confocal microscopy. Animals that did not display clear fluorescent signal (mCitrine for KORD injected animals and GFP for the remaining animals) in the injected cortical area were excluded from the analysis. Immunohistochemistry was performed on a subset of brains to determine the extent of KORD expression.

#### *In vitro* electrophysiology

Neonatal mice were anaesthetized with 4% isoflurane (in O_2_) prior to decapitation followed by rapid dissection of the cerebral cortex in cold, carbogenated (95% O_2_/5% CO_2_) cutting artificial cerebrospinal fluid (ACSF) of the following composition (in mM): 72 sucrose, 83 NaCl, 2.5 KCl, 26 NaHCO_3_, 1 NaH_2_PO_4_, 3.3 MgSO_4_, 0.5 CaCl_2_, 10 glucose (300-310 mOsm; all chemicals were purchased from Sigma unless otherwise stated). Coronal cortical slices were cut in cold cutting ACSF using a vibratome (VT1200, Leica) and allowed to recover in cutting ACSF maintained at 34°C for 30 min and then at room temperature (RT) for a minimum of 30 min. Slices selected for recording were moved to a recording chamber containing RT ACSF of the following composition (in mM): 119 NaCl, 2.5 KCl, 26 NaHCO_3_, 1.3 NaH_2_PO_4_, 1.3 MgSO_4_, 2.5 CaCl_2_, 10 glucose (300-310 mOsm). Individual neurons were visualized using infrared (IR)-differential interference contrast (DIC) microscopy under a 40X objective. Layer location was confirmed *post hoc* under 4X or 10X magnification. Whole-cell patch-clamp recordings were performed using a Multiclamp 700B amplifier and a Digidata 1440A digitizer (Molecular Devices, US). Target neurons were patched with borosilicate glass micropipettes (Harvard Apparatus, UK; 6-9 MΩ resistance), forged using a PC-100 puller (Narishige, Japan). For GABAergic connectivity experiments, recording pipettes were filled with a Cs-based intracellular solution of the following composition (in mM): 100 gluconic acid, 0.2 EGTA, 5 MgCl_2_, 40 HEPES, 2 Mg-ATP, 0.3 Li-GTP (pH 7.2 using CsOH; 270-280 mOsm). When recording intrinsic properties or glutamatergic connectivity, pipettes were filled with a K-based intracellular solution of the following composition (in mM): 128 KGluconate, 10 HEPES, 4 NaCl, 5 Mg-ATP, 1 Li-GTP, 0.5 CaCl_2_ (pH 7.2 using KOH; 270-280 mOsm). For loose-seal cell-attached recordings, pipettes were filled with recording ACSF. The internal solution was supplemented with Biocytin (0.3%) to enable morphological reconstruction of the recorded neurons. All recordings were sampled at 20 kHz. Access resistance (Ra) was monitored during recording and not compensated; recordings were discarded if Ra increased more than 20% of its initial value. Intrinsic electrophysiological properties were determined in current-clamp configuration using negative and positive current injection steps (500 ms) of increasing amplitude. Intrinsic electrophysiological data were analyzed through a custom-made script written in Python (Jupyter Lab v1.0.2) to extract 11 electrophysiological features, with similar methods to Scala et al. (2019).

Laser-scanning photostimulation (LSPS) was performed as previously described [8, 9]. Prior to photostimulation, slices were maintained for a minimum of 6 min in high-divalent cation (HDC) ACSF of the same composition of the recording ACSF but with higher concentration (4 mM) of MgCl_2_ and CaCl_2_ and MNI-caged glutamate (200 µM; Tocris Bioscience, UK) or DMNPE-caged ATP (200 µM; Tocris Bioscience). UV laser pulses (355 nm; DPSL-355/30, Rapp OptoElectronic, Germany) were directed to the slices through a galvanometer scanner system (UGA-42; Rapp OptoElectronic) and focused through a 10X objective. LSPS grid was characterized by 11x17 spots stimulated in a pseudorandom sequence and spaced ∼50 µm; long duration (100 ms) laser pulses were administered at 1-2 Hz frequency. To record the whole extent of the cortical thickness, LSPS grid was moved and for each grid position a minimum of 3 sequences were recorded in voltage clamp configuration. To assess GABAergic connectivity, laser-evoked inhibitory postsynaptic currents were recorded by clamping the cell at the reversal potential for glutamate (E_Glut_) determined empirically. For glutamatergic connectivity experiments, laser-evoked excitatory postsynaptic currents (EPSCs) were recorded by clamping the cell near RMP (Vh = −60 mV).

Analysis of LSPS data was performed with a customized MATLAB (R2020a, MathWorks) toolbox. Analysis of this dataset requires the definition of a putative window for monosynaptic connectivity following the laser stimulus. For glutamatergic connectivity mapping, this window was set according to previously published data, obtained in mice at comparable ages (40 – 337 ms) [8]. For GABAergic connectivity mapping, control experiments were performed to determine the putative monosynaptic window for analysis as well as to calibrate the power of the laser used for further experiments. Lhx6-eGFP+ interneurons in layers 2-5 (n = 19) of visual and somatosensory cortices were targeted for loose-seal cell-attached electrophysiological recordings. Application of the standard LSPS protocol at multiple laser power levels triggered spiking activity in the recorded cell used. Optimal laser power was selected as the one capable of triggering robust firing of the cell when the laser was fired at the soma but weak or no spiking activity when the laser was fired at adjacent locations. Postsynaptic current Within this monosynaptic window, whenever the current traces crossed a threshold set at 6 times the SD of the baseline current, the area under the curve (AUC) was measured as a proxy of the strength of the connectivity. For glutamatergic maps, all EPSCs in the trace were identified with the template search event detection algorithm implemented in Clampfit (v10.7, Molecular Devices). Only EPSCs falling within the monosynaptic window were selected and their amplitude summed in each pixel of the LSPS grid. Multiple maps were then averaged, layer boundaries were manually determined using the DIC image acquired under 10X magnification and the final heatmap was generated. Normalized connectivity maps were obtained by dividing value in each spot by the sum of all pixels. Columnar profiles were obtained by summing all values for each line in individual heatmaps; layer distribution of inputs was calculated by summing all inputs occurring in all pixels belonging to a given cortical layer. Total inputs were obtained by summing values in each pixel of the map. Average maps per groups were obtained by aligning each individual map to the L4-L5 boundary and averaging corresponding pixel values.

#### *In vitro* optogenetics

*In vitro* optogenetics experiments were performed using a 470 nm light emitting diode (LED; Thorlabs Inc., US or CoolLed, UK); focused onto the recorded neuron using a 40X objective. Light-evoked IPSC were recorded voltage-clamping the target cell near the reversal potential for glutamate (Vh = 0 mV) whereas EPSCs were recorded near RMP (Vh= −60 mV). Brief (1 or 10 ms) LED pulses were used to trigger synaptic events in the recorded cell; an inter-stimulus interval of 20-50 s was used to allow for channelrhodopsin-2 (ChR2) recovery. Different LED power intensities were employed to determine power-dependent responses of recorded neurons. When studying GABAergic connectivity, a minimum of 6 trials were recorded for each LED power intensity value. For thalamic input experiments, the minimal stimulation power value was found by increasing the LED power in consecutive sweeps; a minimum of 6 trials were then recorded at the minimal stimulation power level. Analysis of *in vitro* optogenetics data was carried out with customized scripts in Python. Light-evoked synaptic events were identified when the LED stimulus induced a deflection in current larger than 9 times the SD of the baseline. A large detection window (0 – 50 ms from LED stimulus onset) was used to account for possible developmental effects on ChR2. For events within the monosynaptic detection window, synaptic event features were measured, such as peak amplitude, onset latency (obtained by linear regression fit between the value at 20 and 80% of the peak amplitude), onset jitter (standard deviation of the latency), and decay time constant (tau; monoexponential fit to the decay phase of the synaptic event). Percentage of synaptic events occurrence was calculated throughout all trials at each combination of LED power intensity or minimal stimulation LED power. A neuron was considered responsive if synaptic events occurred in at least 33% of the trials at maximal (or minimal for thalamic input experiments) LED power level. For thalamocortical input experiments, relative latency for cells other than L4 pyramidal neurons was calculated by subtracting the latency of a synaptic event to the average obtained from a L4 pyramidal cells patched within the same slice. In a subset of experiments, the following chemical compounds were used to affect optogenetically-evoked synaptic events (all purchased from Tocris Bioscience): Bicuculline methiodide (10 µM), Tetrodotoxin citrate (TTX, 1 µM) and 4-aminopyridine (4-AP, 50 µM). Drugs were incubated for a minimum of 10 min before applying the optogenetic protocol.

#### Morphological reconstruction

Following electrophysiological experiments, slices containing biocytin-filled cells were fixed in 4% paraformaldehyde (PFA; in phosphate-buffered saline, PBS) overnight at 4°C. Slices were then rinsed in PBS and incubated into 0.05% PBST containing Streptavidin-Alexa568 or Streptavidin-Alexa488 (1:500, Molecular Probes, US) for 48-72h at 4°C. Finally, slices were washed in PBS and mounted on histology slides with Fluoromount (Sigma) mounting medium. Slices were imaged through an Olympus FV1200 confocal microscope equipped with a 10X dry objective. Z-stack images were acquired to maximize imaging of all neuronal processes to allow offline morphological reconstruction. Image analysis was performed with Fiji-ImageJ software (NIH, US): morphological reconstruction was manually performed using the Simple Neurite Tracer plugin [10]. Axonal and dendritic arbours were reconstructed separately; throughout the experimental Chapters, pyramidal neurons morphology reconstructions show only the dendritic arbour whereas interneuron morphology reconstructions show the axon.

**Figure S1.**
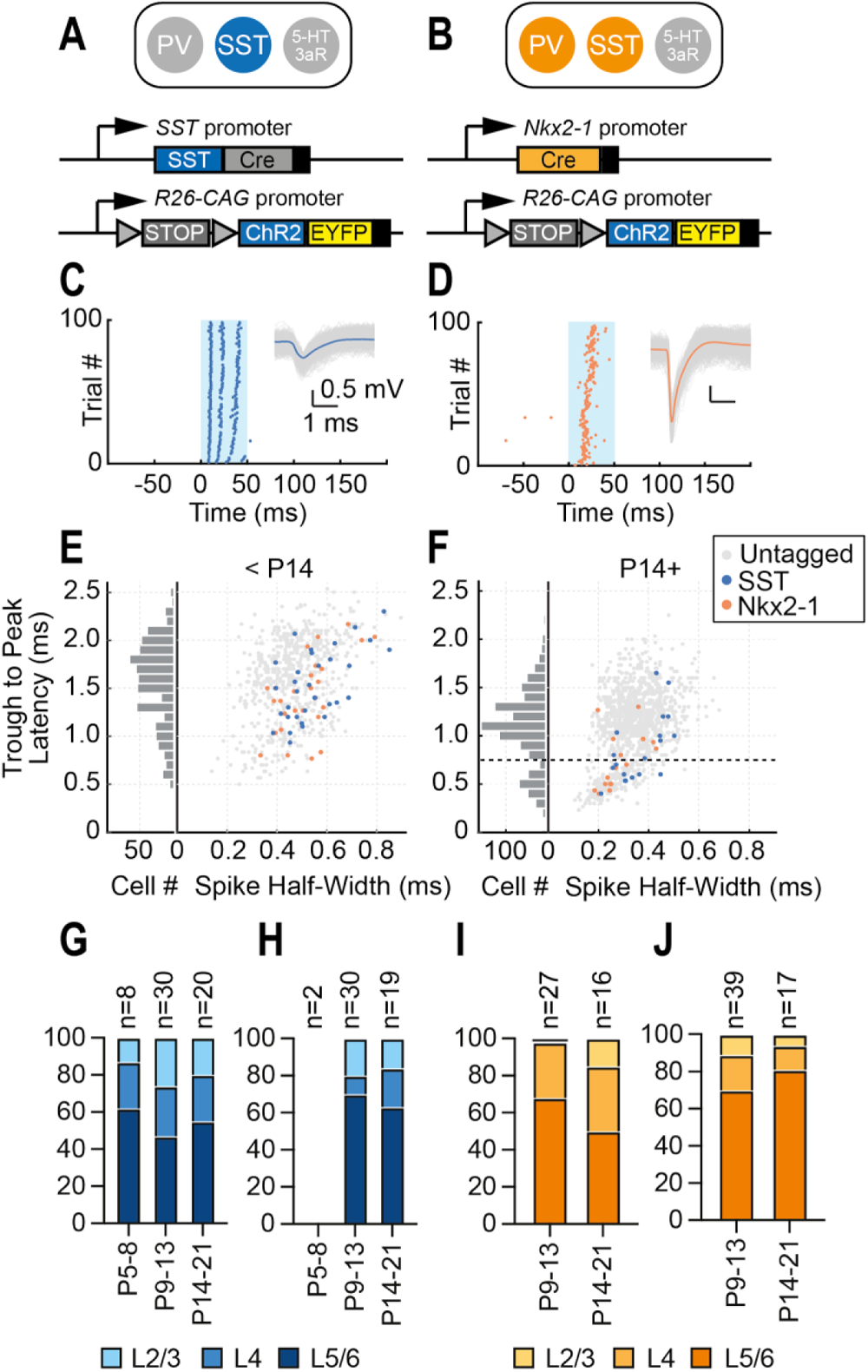
Conditional expression of ChR2 permits optotagging of postnatal GABAergic interneurons across the depth of S1BF and V1 in postnatal neocortex. **(A**) Genetic strategy to conditionally label the SST subtype of GABAergic interneuron. (**B**) Corresponding approach to get both PV and SST interneuron using the *Nkx2-1Cre* line. (**C**) Raster plot of optotagged P12 SST interneuron with spikes elicited during 50 ms 470 nm LED flash (light blue box). Inset, spike waveforms for the single unit; grey lines indicate individual spike waveforms, blue line shows average spike waveform. (**D**) Corresponding data for P13 Nkx2-1 SU. (**E**) Scatter plot of spike half-width *vs.* spike through to peak latency (right) and histogram plot for spike through to peak latency (left) for SUs obtained from animals before eye opening. Grey dots, untagged SUs, blue dots, optotagged SST interneurons, orange dots, optotagged Nkx2-1 interneurons. (**F**) Corresponding data for SU recorded after eye opening. Horizontal dashed line, arbitrary cut-off spike through to peak latency value (0.75 ms) used to identify fast-spiking (FS, below the line) *vs*. regular spiking (RS, above the line) SU. (**G**) Proportion of SST single units recorded per layer through development in S1BF. (**H**) Corresponding data for V1. (**I)** Proportion of Nkx2-1 SU recorded per layer through development in S1BF. (**J**) Corresponding data for V1.

**Figure S2.**
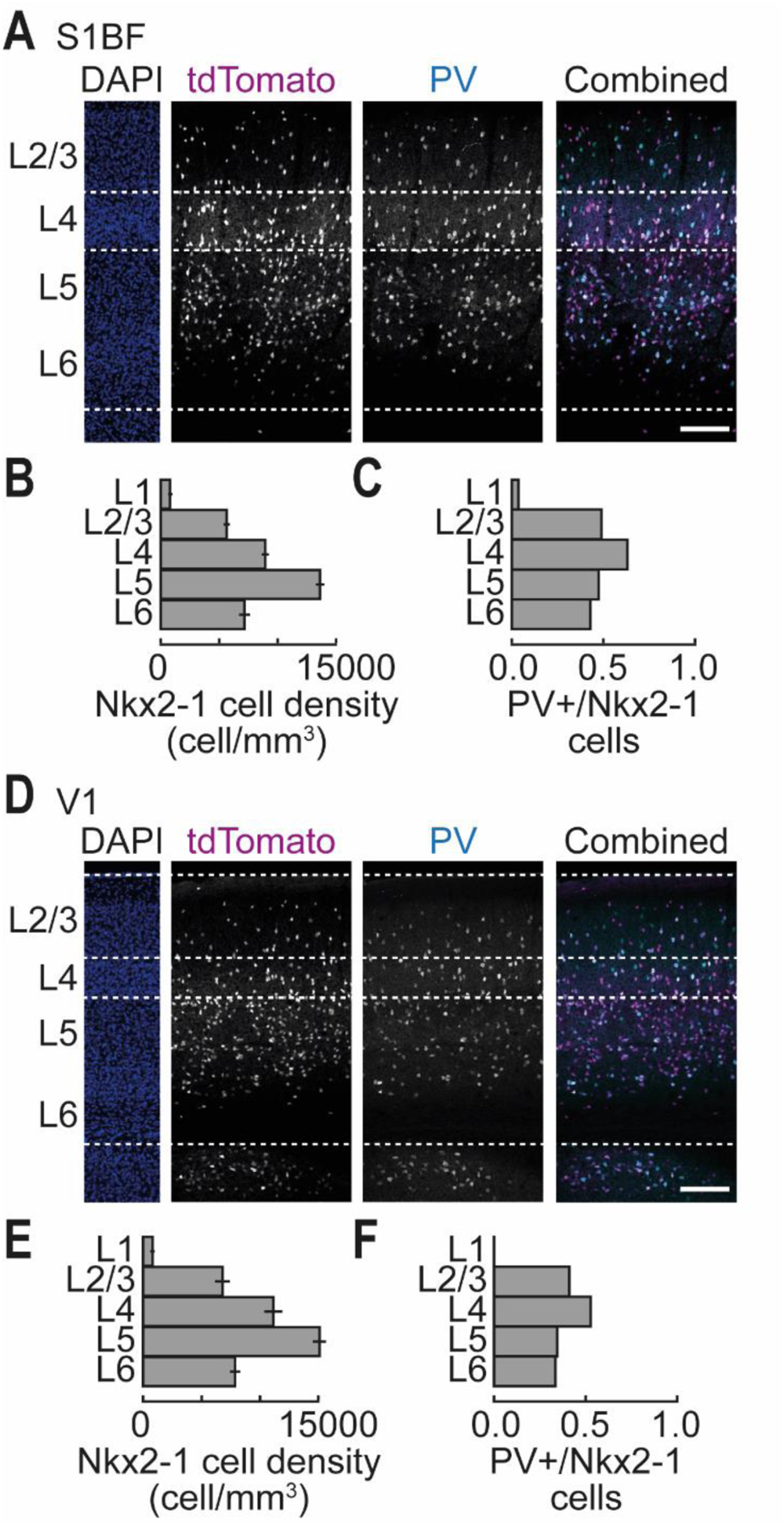
The *Nkx2-1Cre* line labels similar proportions of PV interneurons across both S1BF and V1 in postnatal mouse neocortex. (**A**) Immunohistochemistry for tdTomato and the calcium binding protein, parvalbumin (PV), in somatosensory cortex (S1BF) of P21 mouse *Nkx2-1Cre;Ai9* pup. (**B**) density of Nkx2-1 interneurons across layers (±SEM); Layer location is statistically significant with regards to the cell density data (Two-way ANOVA: F (4,64) = 160.7, p ≤ 0.001). (**C**) Proportion of Nkx2-1 cells that are PV+ across the layers in S1BF. (**D-F**) Corresponding data for P21 V1. (**E**) Layer location is significant for cell density (Two-way ANOVA: F (4,64) = 104.6, p ≤ 0.001). Scale bar: 100 μm.

**Figure S3.**
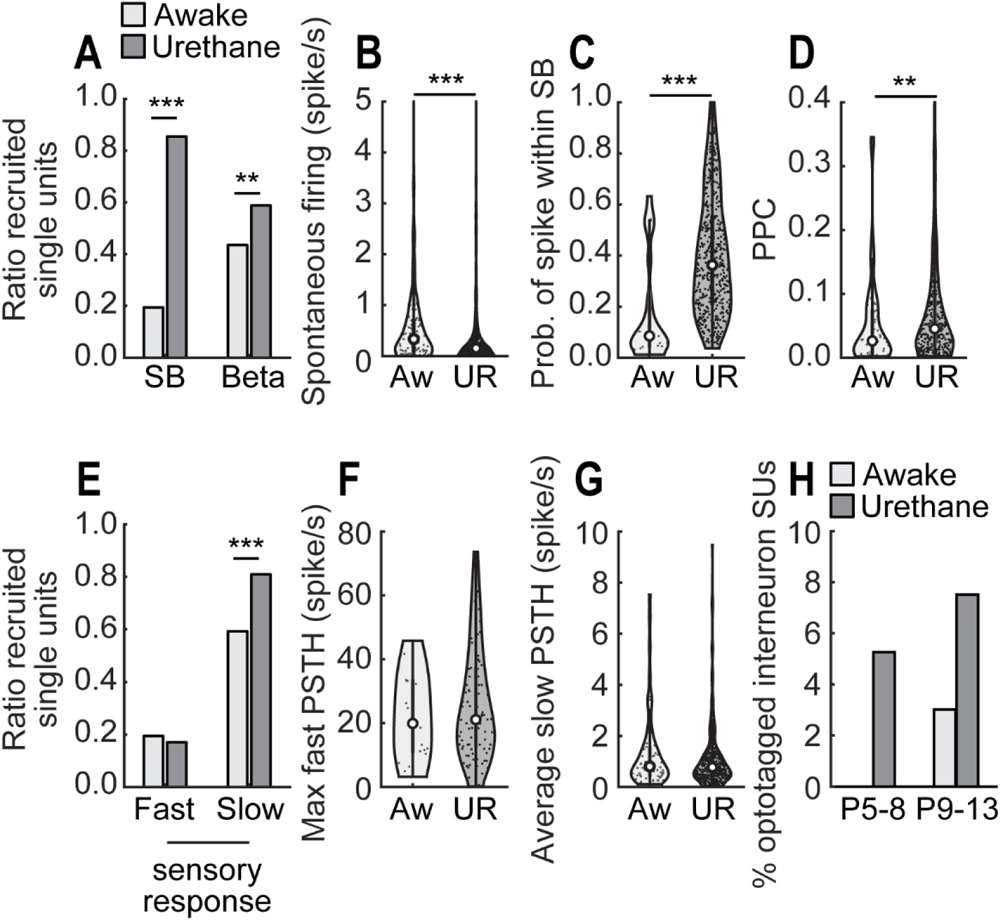
Comparison of spontaneous and sensory evoked activity of RS single units recorded in awake (Aw) and urethane (UR)-anaesthetized mice prior to active sensation (<P14) (A) Bar plot of the ratio of RS SUs significantly entrained by spindle bursts (Chi-square test: Χ^2^ = 214; p ≤ 0.001) or by β frequency oscillations (Chi-square test: Χ^2^ = 8.7; p =-0.003) obtained from Aw head-fixed (light grey) vs. UR-anaesthetized (dark grey) mice. (B) Violin plot showing a difference in spontaneous firing rate of RS SUs in Aw vs. UR mice (Wilcoxon rank sum test: Z = -3.7, p ≤ 0.001). (C) The probability of RS units spiking within spindle bursts (SB) is increasing by UR (Wilcoxon rank sum test: Z = 4.6, p ≤ 0.001) with (D) an increase in pairwise phase consistency (PPC) value between RS units spiking and β-frequency oscillations (Wilcoxon rank sum test: Z = 2.85, p = 0.004). (E) Ratio of single units responsive during the fast (Chi-square test: Χ^2^ = 0.36; p = 0.5) and slow (Chi-square test: Χ^2^ = 24.7; p ≤ 0.001) phases of the postnatal (P9-13) visual responses. The maximum firing rate of responsive RS SUs during (F) the fast phase of the visual response was not altered by UR (Unpaired t-test: t(121) = 1.03, p = 0.3), nor during (G) the slow phase of the visual response (Wilcoxon rank sum test: Z = 0.3, p = 0.7). (H) Proportion of optotagged interneuron single units identified during early postnatal development.

**Figure S4.**
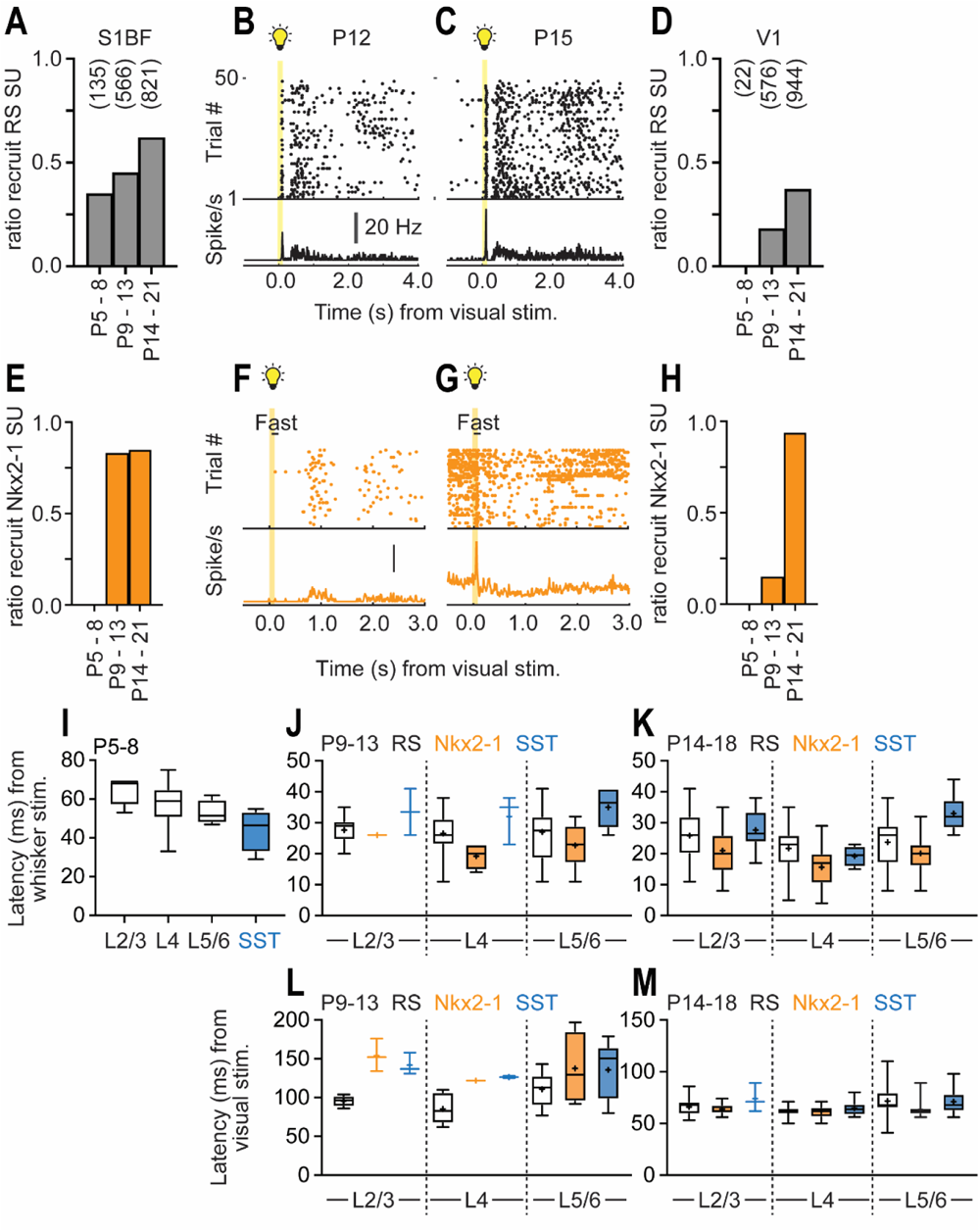
Differences in the recruitment of regular spiking (RS) and Nkx2-1 single units during short latency sensory response over postnatal development. (A) Proportion of RS recruited in S1BF over postnatal development; total number of units identified at each time point shown in parentheses. (B) Representative raster plot of RS unit recruited in response to a flash of light prior to eye opening; onset and duration of stimulus indicated by the yellow bar and light bulb. (C) Corresponding data for an RS single unit record post-eye opening. (D) Proportion of RS units exhibiting a short latency response in V1 over development. (E-H) Corresponding data for optotagged Nkx2-1 single units. Average latency (ms) from multi-whisker stimulation to recruitment of RS (white), Nkx2-1 (orange) and SST (blue) single units in S1BF at (I) P5-8, (J) P9-13), and (K) P14-18. (L, M) Corresponding data for single units recorded in V1 at (L) P9-13 and (M) P14-18).

**Figure S5.**
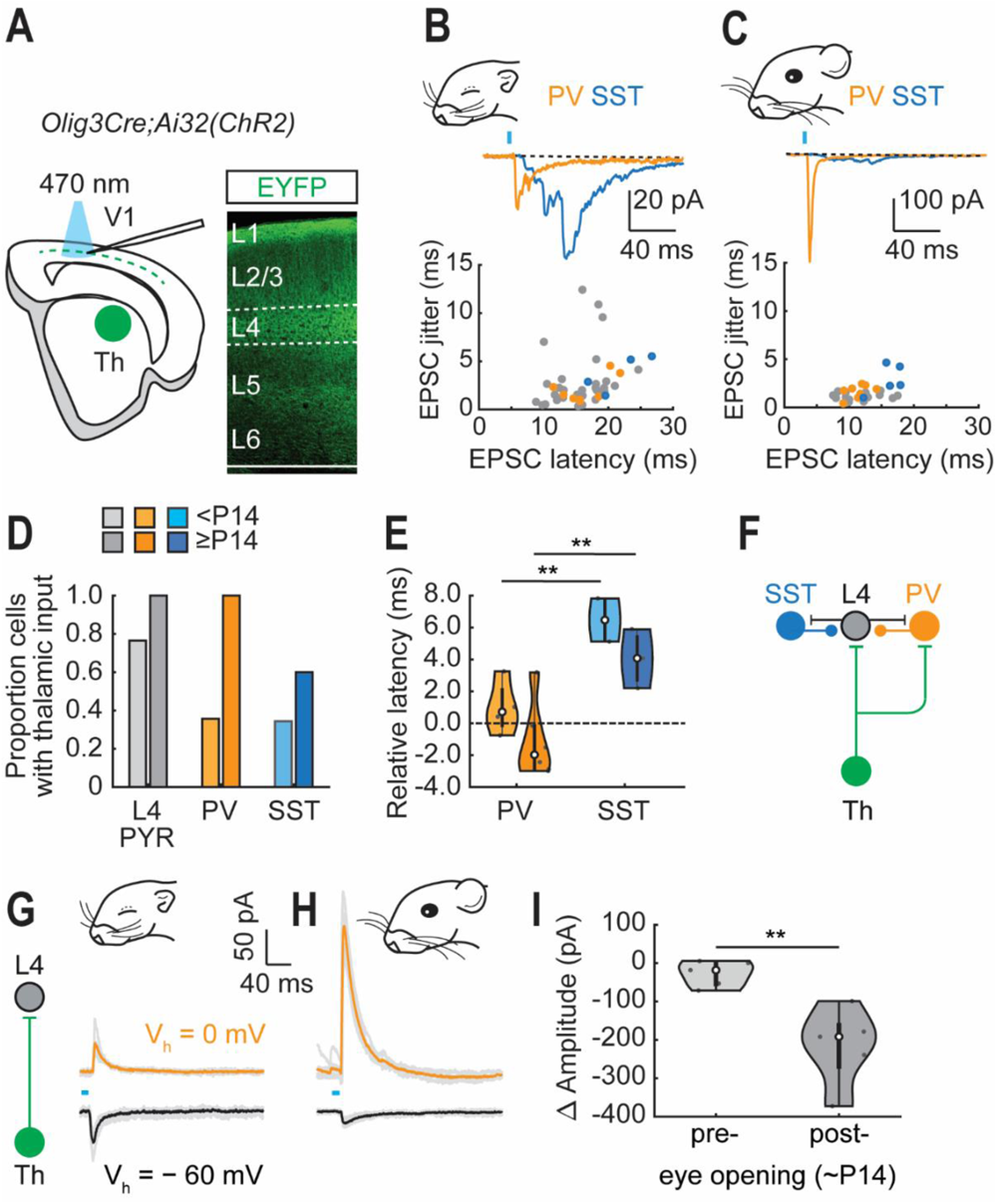
PV but not SST interneurons receive short latency thalamic input with feed-forward inhibition present in V1 following eye-opening. (**A**) Experimental set-up for investigating thalamic innervation in acute *in vitro* slices of postnatal V1. (**B**) Plot showing latency and jitter (SD of onset) of 470 nm light-evoked EPSCs recorded prior to eye opening. PYR cells (grey), PV (orange) and SST (blue) interneurons; inset, representative traces from PV and SST interneurons. (**C**) Corresponding data obtained from recording performed post-eye opening. (**D**) Proportion of cells with short latency (20 / 18 ms), low jitter (3.5 / 2.5 ms), putative monosynaptic thalamocortical (TC-)EPSCs. (**E**) Relative latency of onset TC-EPSCs in PV and SST interneurons compared to L4 PYR cells; color scheme as in panel (D). (**F**) Schematic of the early postnatal thalamocortical circuit within L4 V1. Development of feed-forward inhibition onto L4 glutamatergic neuron in V1 through optogenetic stimulation of ChR2-expressing thalamic terminals (**G**) prior to eye opening (P9-13) and (**H**) post-eye opening (P14-18). (I) The amplitude of light-evoked EPSC minus that of the IPSC recorded from the same cell, pre-and post-eye opening (unpaired t-test: t(8) = -3.9, **p ≤ 0.01).

**Figure S6.**
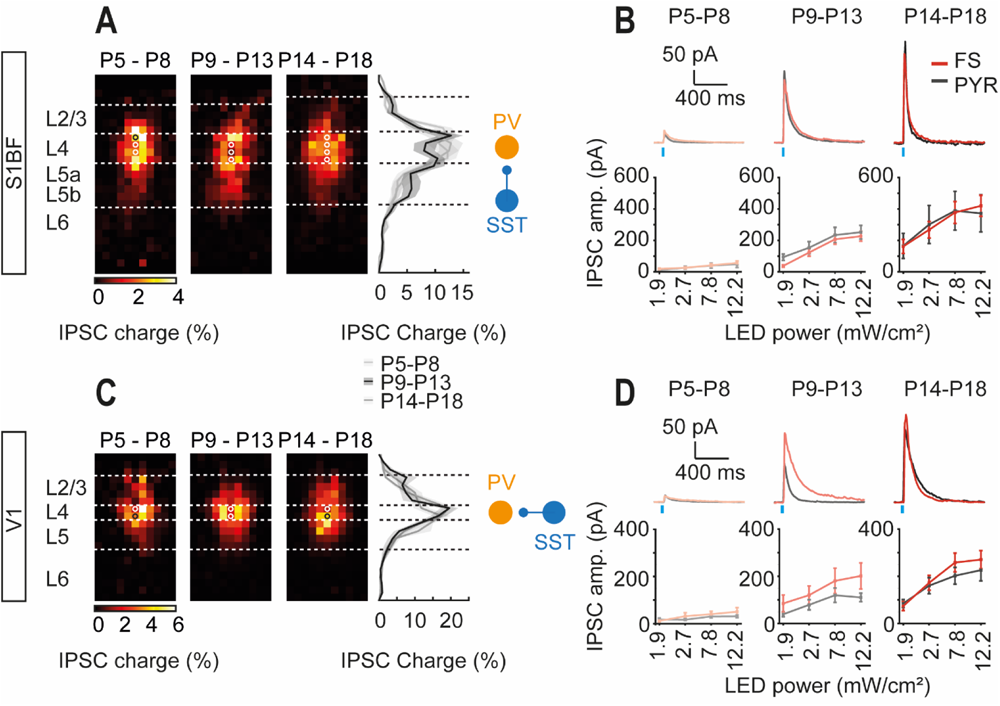
A prominent disinhibitory connection mediated by SST onto PV interneurons emerges prior to active sensory exploration in both S1BF and V1. (A) Total GABAergic input onto L4 PV interneurons across postnatal development in S1BF mapped using LSPS; right panel, plot of normalized input profiles revealing increase L5 input onto L4 PV+ interneurons at P9-13. (**B**) IPSCs recorded in PV interneurons (red traces) and Pyramidal cells (PYR)(black traces) in response to optogenetic stimulation of SST interneurons over development; bottom plots, IPSC amplitude (amp.) observed across increasing LED power (mW/cm^2^). (**C-D**) Corresponding data for V1 L4 PV interneurons.

**Figure S7.**
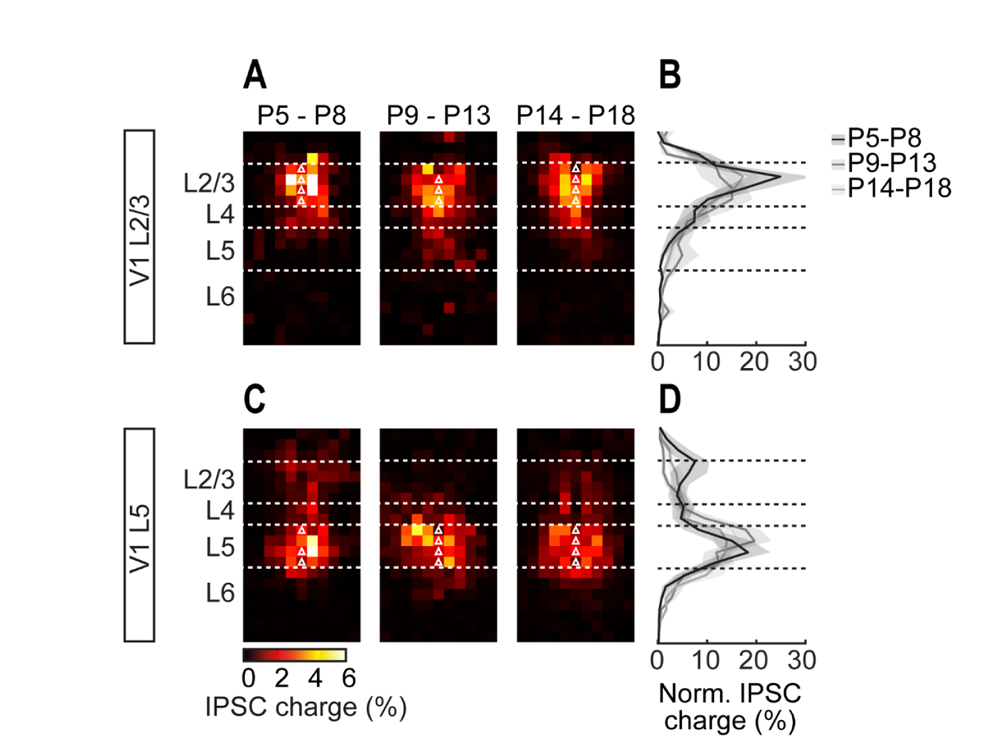
The extent of local versus translaminar GABAergic inputs varies by cortical layer and timepoint in the development in postnatal V1. (A) Total GABAergic into L2/3 pyramidal cells in V1 recorded between P5 and P18. (B) Normalized input profile for the cells shown in (A) showing predominantly local input. (**C-D**) Corresponding data for L5 pyramidal cells in V1.

**Figure S8.**
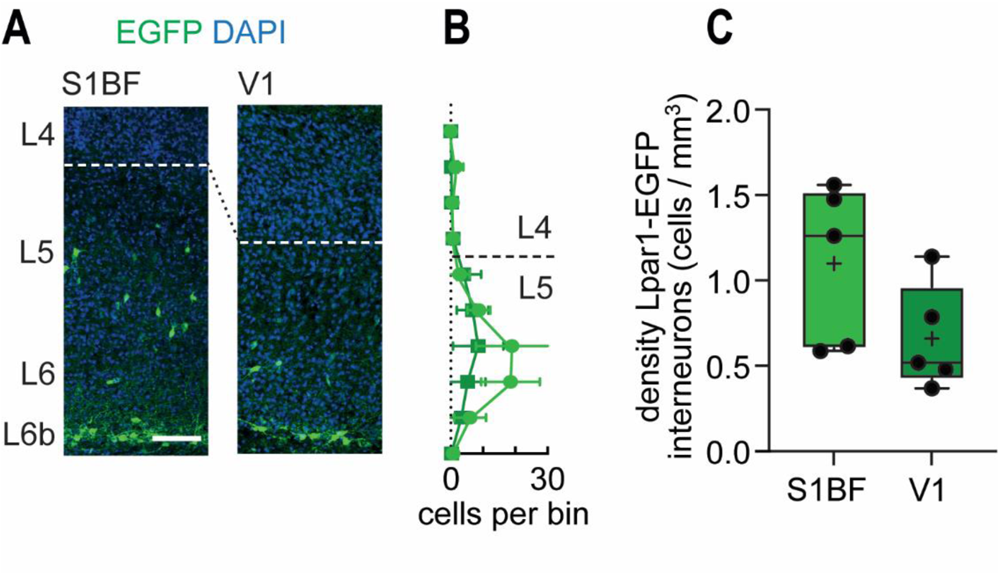
Lpar1-EGFP interneurons are found at similar density across S1BF and V1 in postnatal mouse neocortex. (**A**) representative images of GFP cells, both infragranular interneurons and subplate cells in P7 S1BF (left) and V1 (right). (**B**) Total number of EGFP-positive neurons, excluding subplate, measured across the normalized depth of S1BF (light green)(counts from n=5 brains from separate litters) and V1 (dark green)(n=5 brain). (**C**) density of Lpar1-EGFP cells in both cortical areas (unpaired t-test: t(18) = 1.469, P = 0.159).

**Figure S9.**
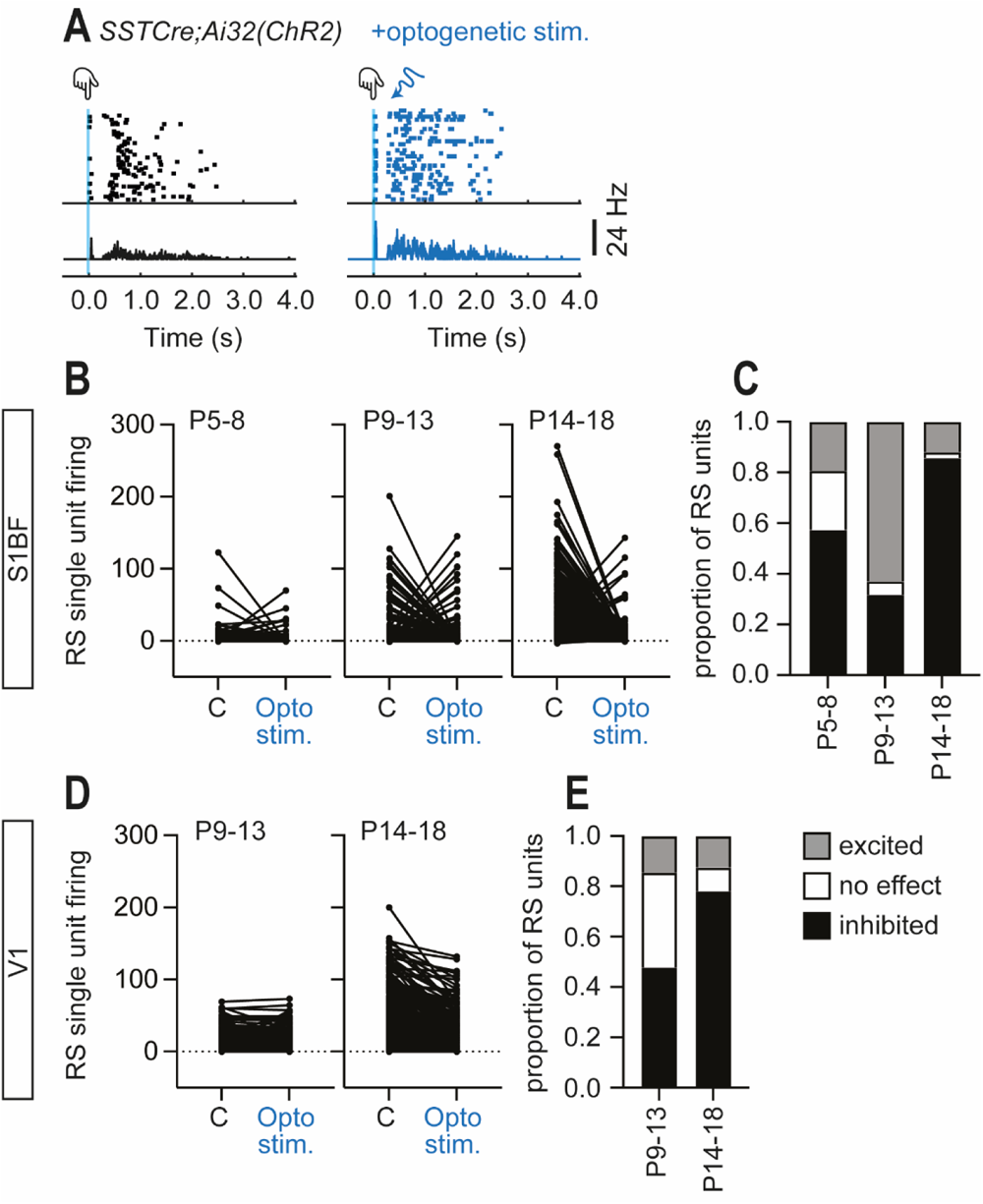
Optogenetic activation of SST interneurons alters single unit sensory evoked responses in a varied manner in both postnatal S1BF and V1. (**A**) Representative example showing the effect of optogenetic stimulation of SST interneurons coincident with multi-whisker stimulation in S1BF at P8. Left black trace, control; right, blue trace, with SST interneuron optogenetic stimulation. (**B**) Effect of optogenetic stimulation of SST interneurons on RS pyramidal cells in S1BF across the developmental time windows studied. (**C**) summary data showing the proportion of excited (grey), neutral (white) and inhibited (black) RS units. (D-E) Corresponding summary data for V1.

**Figure S10.**
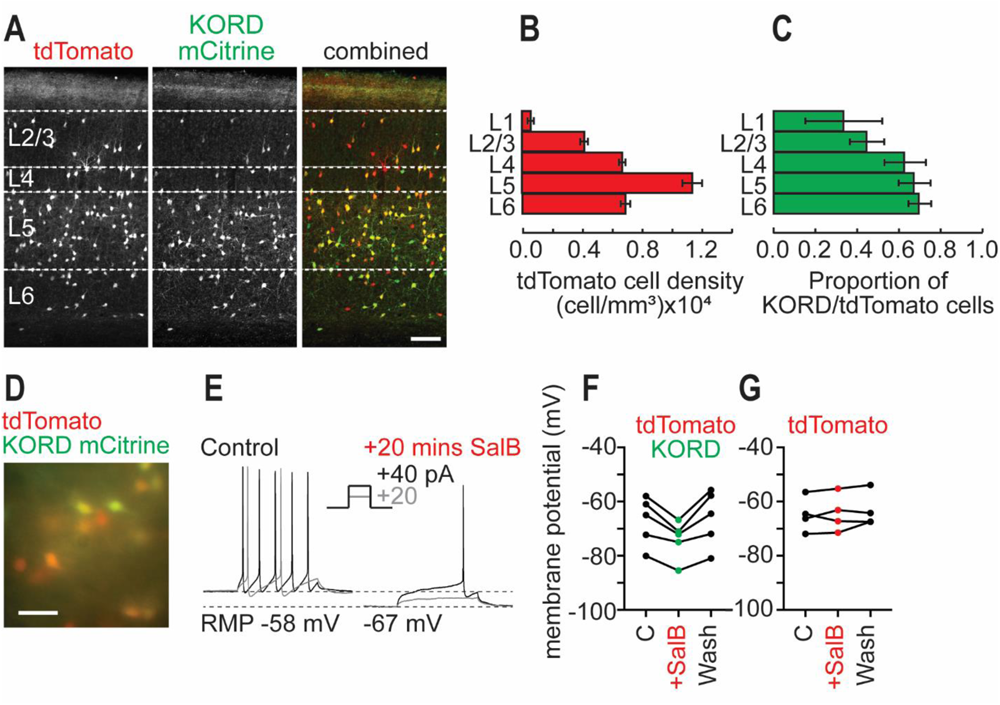
AAV8-KORD-ires-MCitrine transduction of SST interneuron in postnatal sensory cortices leads to SalB-controlled changes to SST interneuron excitability. (A) Representative images of KORD mCitrine transduction in a tdTomato labelled SST interneurons in P11 V1. (B) Quantification of SST interneuron density and (C) proportion of KORD-expressing/tdTomato SST interneurons across the layers of sensory cortex (n = 6 animals). (D) Visualizations of tdTomato and KORD mCitrine cells in acute *in vitro* slices. (E) Representative current clamp traces showing the spiking response of a P7 KORD-expressing SST interneuron to 20 (grey trace) and 40 pA (black) in control ACSF and ACSF containing 100 μm SalB. The average membrane potential recorded in (F) KORD-expressing and (G) control tdTomato-only SST interneurons in control (C), application of ACSF + SalB, and following 20 mins. washout of drug (Wash).

**Figure S11.**
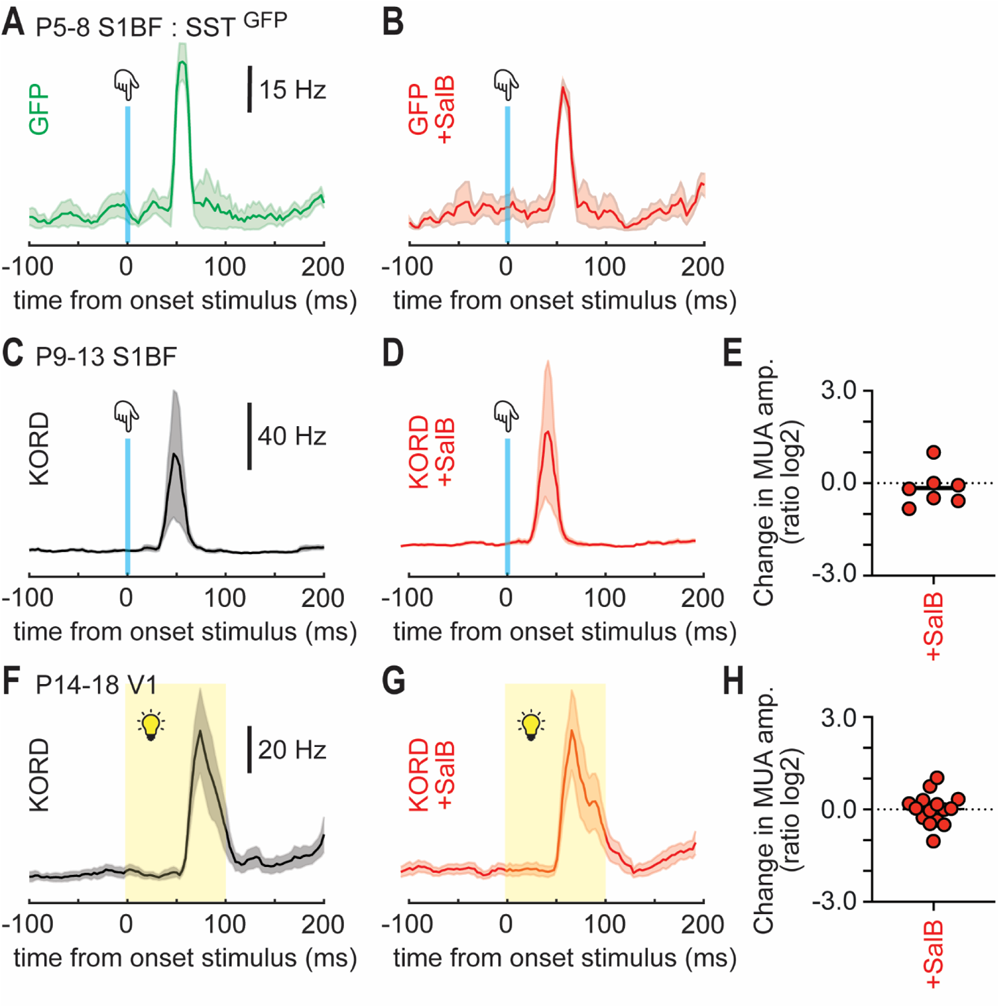
KORD-expression provides a controlled and effective means for regulating interneuron activity at early postnatal ages. **(A)** Baseline MUA recorded in response to whisker stimulation in a postnatal mouse injected with AAV-GFP. (**B**) Addition of the KORD-specific agonist Salvinorin B (SalB)(1 mg/kg; 0.1 mg/ml in saline with 1% DMSO) results in no change in MUA activity. (**C)** Average baseline MUA activity recorded in response to multi-whisker stimulation between P9 and P13. (**D-E**) Addition of SalB had no effect on the either (D-E) the average whisker response between P9-13 nor (**F-H**) the visual response recorded post-eye opening (P14-18).

